# UPLC-MS analysis of characterization of bioactive compounds produced by endophytic *Bacillus tequilensis* ALR-2 from *Aloe vera* plant

**DOI:** 10.1101/2020.07.23.217687

**Authors:** Mushafau Adewale Akinsanya, Joo Kheng Goh, Adeline Su Yien Ting

## Abstract

Endophytic *Bacillus tequilensis* ALR-2 isolated from *Aloe vera* plant, was found to have antimicrobial activities attributed to a series of lipopeptide antibiotics and biosurfactants produced. Partial-purification of ethylacetate extracts by column chromatography revealed that fraction SF5 (5mg mL^−1^) effectively inhibited *Bacillus cereus* and *Staphylococcus aureus* (28.6±0.5 and 30.3±0.5 mm zone of inhibition, respectively), and was significantly more effective than the standard antibiotics Ciprofloxacin (5μg) (23.6 ±0.5 and 18.6 ±0.5 mm, respectively) and Kanamycin (30μg) (20.3 ±0.5 and 13.6 ±0.5 mm, respectively). UPLC-MS analysis of SF5 showed three series of ion peaks. The quadrupole-orthogonal ionization time-of-flight mass spectrometer revealed [M+H]^+^ of 678.4601; 679.4690 and 680.5076 which corresponded to the oligopeptide antibiotic monamycin (mol. wt. 677.84). The [M+H]^+^ of 1119.8870; 1120.8992 and 1121.9119 corresponded to antibiotic complex 61-26 (mol. wt. 1120.348), a lipopeptide. The third peak of [M+H]^+^ of 1356.0151 and 1358.0664 could not be classified to existing known compounds in the literatures.

## Introduction

Endophytes are bacteria or fungi, which reside in the tissues of higher plants without causing any symptoms or apparent harm to the host. These microbes are not pathogenic to the host, neither are they providing benefit to other microbial residents [1]. They have been found to have a wide range of antimicrobial activities, hence are important sources of antimicrobial substances [2]. In recent years, endophytes have been found to produce novel metabolites exhibiting a variety of biological activities against different pathogens [3, 4]. Thus, endophytes have become increasingly popular for isolation and characterization of bioactive compounds [5].

Some endophytes have been found to produce bioactive compounds that may be involved in a symbiotic association with a host plant [6]. In recent studies, it has been recognized that endophytic bacteria play an important role in resistance to diseases and mediate some beneficial role between the endophyte and its host [5, 7]. Similarly, various studies have been conducted on the plant growth-promoting abilities of endophytes and this has indicated their ability to increase plant growth through the improvement of nutrient/mineral recycling, and their ability to inhibit invading phytopathogens [8].

*Bacillus tequilensis* ALR-2 was isolated from *Aloe vera*, along with other endophytic *Proteobacteria, Firmicutes* and *Bacteriodetes*, and was found to produce the most potent antimicrobial bioactive compounds to inhibit gram-positive and gram-negative pathogens [9]. Although studies on *Bacillus* sp. is quite common, there are not many on *Bacillus tequilensis*, particularly on their antimicrobial potential [10]. As such, this study characterized the bioactive compounds from *B. tequilensis* ALR-2 using Ultra-Performance Liquid Chromatography (UPLC) and Mass Spectrometer, which essentially determines accurate mass of compounds. This method is superior due to its efficiency and high resolution of molecular ions compared to other mass-to-charge ratio *(m/z)* analyzers [11]. This study reports on the antimicrobial activities of crude extracts of endophytic *Bacillus tequilensis* ALR-2, and the characterization of the bioactive compounds produced.

## Results and Discussion

*Bacillus tequilensis* ALR-2 is a Gram-positive, endospore-forming, motile and rod-shape bacterium. As reported in our earlier study, the nucleotide sequence of this isolate (16S rDNA) showed the closest similarity to that of *Bacillus tequilensis* (with a homology of 97%) [9]. This strain is reported as an endophyte from *Aloe vera* plant for the first time, with the GenBank accession number KJ689792 assigned [9]. This isolate has excellent antimicrobial activity towards *Staphylococcus aureus, Bacillus cereus, Proteus vulgaris, Klebsiella pneumoniae, Escherichia coli, Streptococcus pyogenes* and *Candida albicans* as documented in our earlier studies [9].

### Antimicrobial activities screening of extracts via TLC-bioautography

The results indicated that only ethylacetate-derived extracts showed bioactivities in the TLC-bioautography (Fig. 1). This suggested the bioactive components are present in the more polar solvent (ethylacetate with polarity index 4.4) than the less and non-polar solvents such as diethylether and n-hexane, respectively. The ethylacetate extract, when subjected to TLC, indicated 2 separate fractions in the chromatogram: A2-1 and A2-2 with Rf values 0.98 and 0.86, respectively (Fig. 1). The two fractions showed antimicrobial activities against *S. aureus* and *B. cereus* (Fig. 1) as the zone of inhibition was wider and appeared to map the two fractions with Rf values (0.98 and 0.86). Hence, the TLC-bioautography technique has enabled us to rapidly identify the fractions with bioactive compounds of interest and the right separating mixture for the isolation of the bioactive compounds [16]. Therefore, *B. tequilensis* (ALR-2) may be said to possess very promising antimicrobial compounds as revealed by the prominent zone of inhibition seen on the chromatogram.

**Figure 1.**
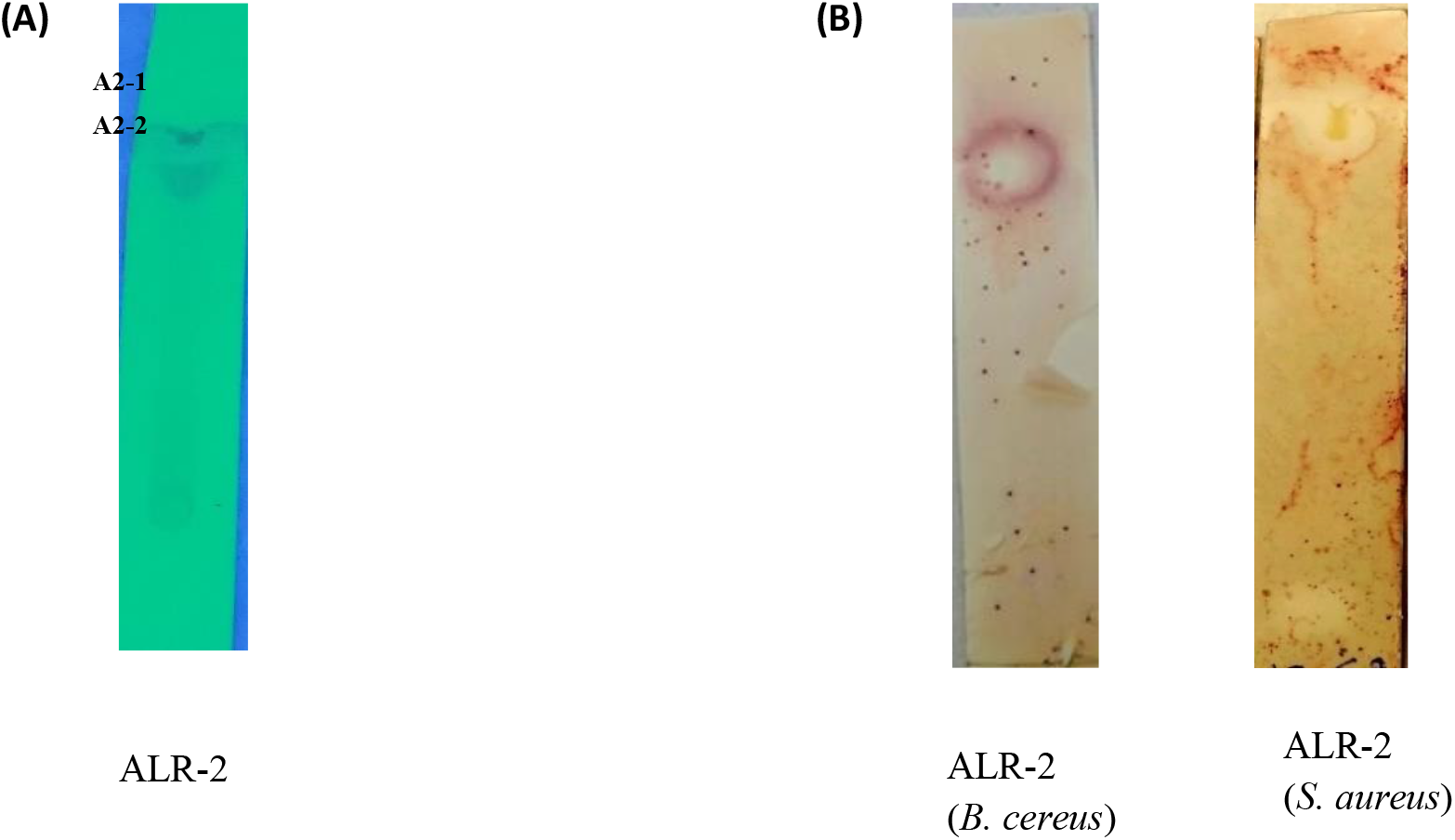
(A) Bioautography plates. Chromatograms plate of ethylacetate extracts of ALR-2 showing fractions (A2-1, A2-2). (B) Clear areas showing inhibition zones of chromatograms when immersed in molten agar seeded with bacterial pathogens, incubated at 37 ± 2°C for 18 h then sprayed with tetrazolium compound and then incubated at 37 ± 2°C for 24 h. (Images scale x2)

### Extraction and purification of bioactive compound

The ethyl acetate extraction of *Bacillus tequilensis* ALR-2 yielded 3.4 mg of lyophilized powder forms (from 1 L broth culture), the highest yield compared to diethylether (2.02 mg) and n-hexane (0.69 mg). Upon fractionation via the LiChroprep RP-18 (15-25 μm) silica gel column (Merck, EMD Millipore Corporation, USA), 10 fractions were obtained (see supplementary Table 1). Fractions F6 and F7 have the highest yield with 6443 and 6283 mg mL^-1^, respectively, while fractions F2 and F3 produce the lowest yield of 110 and 156 mg mL^-1^, respectively. Since ALR-2 was extracted with polar solvent, it is expected to contain more polar compounds than non-polar compounds. The combined three fractions of F7, F8 and F9 gave six subfractions (see supplementary Table 1).

One of these compounds was detected in SF3 and SF6 at retention time 22.9 min with 25.6% and 2.0%, respectively. It was also observed that SF3 was equally active against *S. aureus, B. cereus* and *K. pneumonia*, with zone of inhibitions 9.3 ±0.5, 13.6 ±0.5 and 10.6 ±0.5 mm, respectively (Table 2). We also observed SF6 was active against *S. aureus* and *B. cereus*, and this may be attributed to the compound detected at peak 24.4-minute retention time of 26%. This compound was also detected in SF3 (8%), SF4 (7%) and SF5 (2%) retention time of 24.4 min. The results agreed with statements of researchers’ bioprospecting for new antibiotics from either soil microbes or endophytes, that these microbes produce several antimicrobial and antifungal compounds in their culture, and some exist in isoforms [17–19]. Hence, the most potent of the compound was targeted for identification which we believed can be found in SF5.

### Antimicrobial activity

Seven fractions (F4 – F10) inhibited three to five pathogens tested with zone of inhibition from 12.3 to 20.6 ±0.5 mm, while the first three fractions (F1-F3) appeared to have no inhibition against all the pathogens tested (Table 1). Gram-positive bacteria: *S. aureus, B. cereus, E. faecalis* and *S. pyogenes* were more susceptible to fractions F7-F10 while *C. albicans* was only inhibited by F10 with inhibition zone of 12.0 ±0.5 mm. Other studies have also reported that the bioactive compounds produced by *Bacillus* species is active against gram-positive bacteria pathogens [20, 21]. However, it appeared fractions-F7, F8 and F9 relatively inhibited the pathogens more than fractions F4, F5, F6 and F10 (Table 1). Hence, they were considered to have high bioactivity and were selected for further purification analysis.

**Table 1.** Antimicrobial activities of fractions (F1-F10) against bacteria and *C. albicans* pathogens.

| Fraction<br>(20 mg mL<sup>-1</sup>) | *E. faecalis* | *B. cereus* | *S. pyogenes* | *S. aureus* | *K.*<br>*pneumoniae* | *C. albicans* |
| --- | --- | --- | --- | --- | --- | --- |
| F1-F3 | - | - | - | - | - | - |
| F4 | 13.6±0.5<sup>abc</sup> | 16.3±0.5<sup>bc</sup> | - | 18.3±0.5<sup>b</sup> | - | - |
| F5 | 13.6±0.5<sup>abc</sup> | 17.6±0.5<sup>c</sup> | - | 19.3±0.5<sup>bc</sup> | - | - |
| F6 | 13.0±0.5<sup>ab</sup> | 16.3±0.5<sup>bc</sup> | - | 20.3±0.5<sup>c</sup> | - | - |
| F7 | 15.6±0.5<sup>d</sup> | 15.3±0.5<sup>b</sup> | 14.3±0.5<sup>b</sup> | 19.3±0.5<sup>bc</sup> | - | - |
| F8 | 14.6±0.5<sup>cd</sup> | 15.3±0.5<sup>b</sup> | 14.3±0.5<sup>b</sup> | 20.0±0.5<sup>c</sup> | - | - |
| F9 | 14.3±0.5<sup>bcd</sup> | 16.0±0.5<sup>bc</sup> | 15.0±0.5<sup>b</sup> | 20.6±0.5<sup>c</sup> | - | - |
| F10 | 12.3±0.5<sup>a</sup> | 12.6±0.5<sup>a</sup> | 12.3±0.5<sup>a</sup> | 16.3±0.5<sup>a</sup> | - | 12.0±0.5 |

Data are mean ±SD values. One-way ANOVA was used to analyse all data and means were compared using Tukey's Studentized Range Test (HSD<sub>(0.05)</sub>), with differences considered statistically significant when p<0.05 represented by alphabets (a-d). F1-F10 represent the fractions from first fractionation. Pathogens-*Pseudomonas aeruginosa* ATCC 10145, *Salmonella* Typhimurium ATCC 14028, *Proteus vulgaris* ATCC 8427 and *Escherichia coli* ATCC 25922 were resistance to all the fractions. (-) represent no inhibition zone.

**Table 2.**
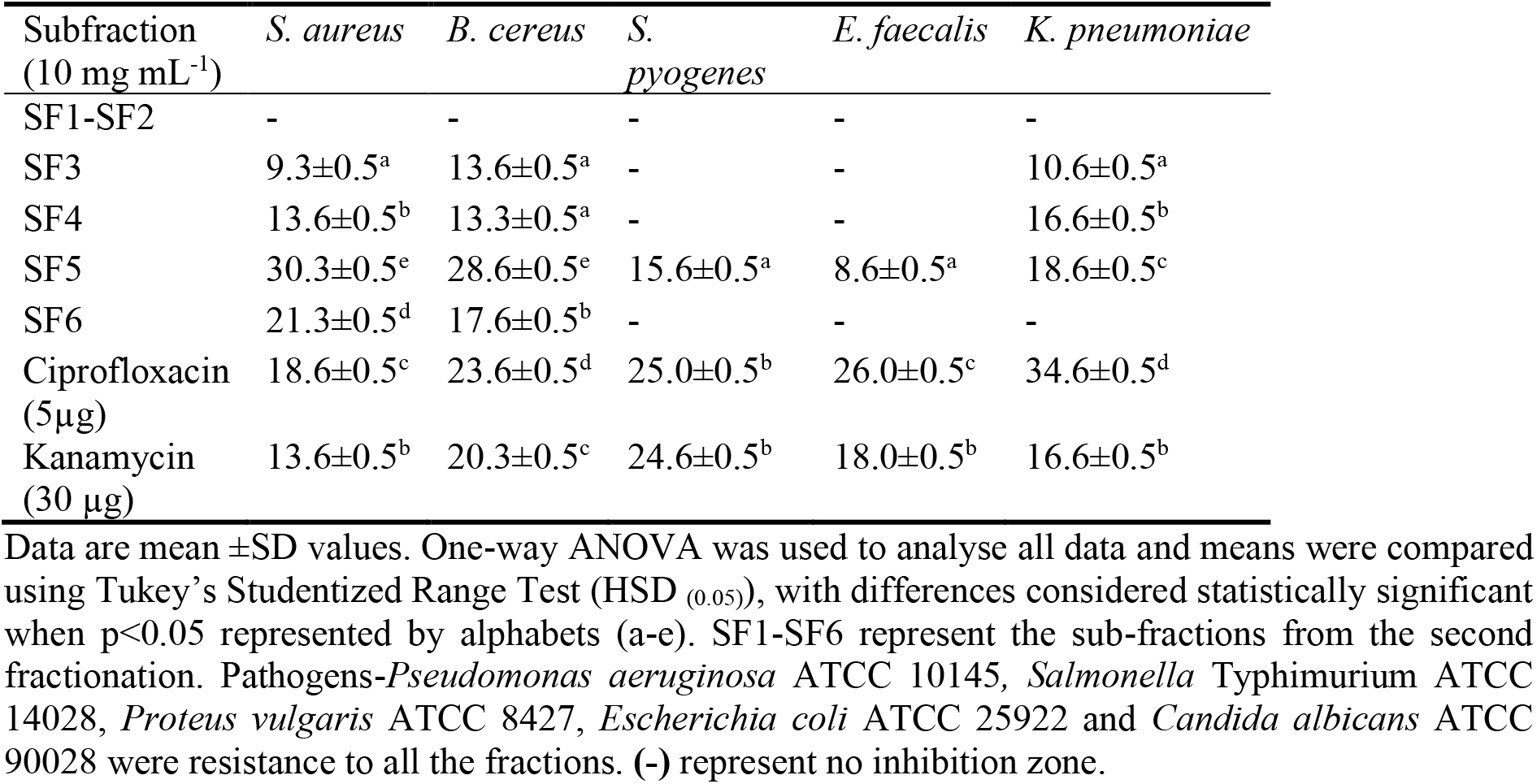
Antimicrobial activities of subfractions (SF1-SF6) against bacteria pathogens.

The antimicrobial activities of the subfractions (SF1-SF6) revealed subfraction SF5 to have the highest antimicrobial activity against all the Gram-positive bacteria (with zone of inhibition from 8.6 to 30.3±0.5 mm) (Table 2). It is interesting to note that antimicrobial activities of SF5 against *S. aureus* and *B. cereus* were significantly higher than the standard antibiotics used; Ciprofloxacin (5 μg) and Kanamycin (30 μg) as positive control (Table 2, Fig. 3). Subfractions SF3 and SF4 also showed antimicrobial activities against three of the five pathogens tested, with zone of inhibition from 9.3 to 16.6 ±0.5 mm. Only subfraction SF6 inhibited two pathogens with zone of inhibition from 10.6 to 21.3 ±0.5 mm. The inhibitory activities can be seen to increase as the major bioactive compound(s) increases in the subfractions. Consequently, this result confirmed the presence of the most active bioactive compound in the subfraction SF5 which has also been revealed by the result of the quantification analysis done with HPLC (Fig. 2).

**Figure 2.**
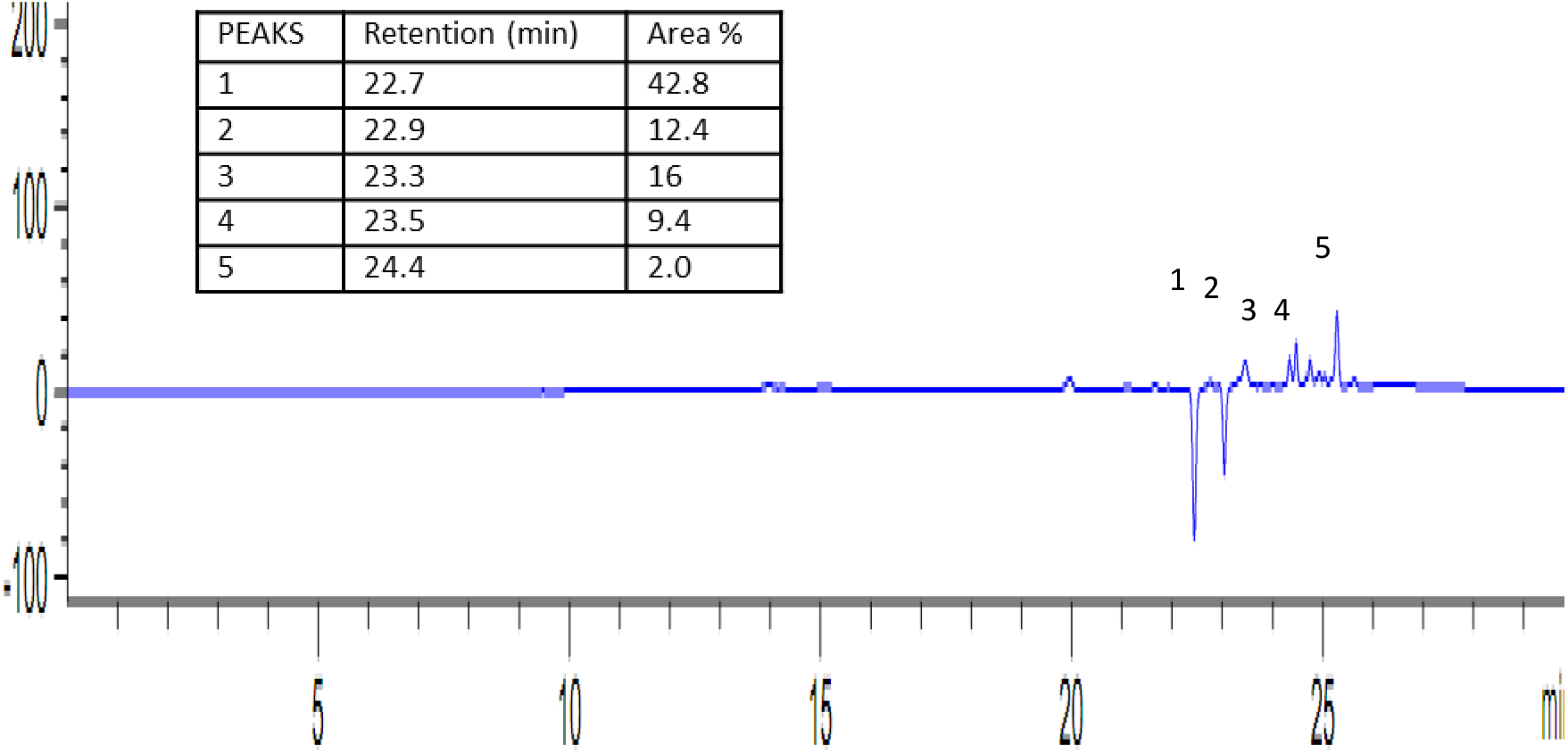
HPLC chromatogram of subfraction SF5 at 293 nm showing five major peaks *(1) = 42.8% area at retention time 22.7 min., (2) = 12.4% at retention time 22.9 min.*, (3) = 16% areas at retention time 23.3 min., (4) = 9.4% area at retention time 23.5 min. *(5) = 2%; 24.4 min.*

**Figure 3.**
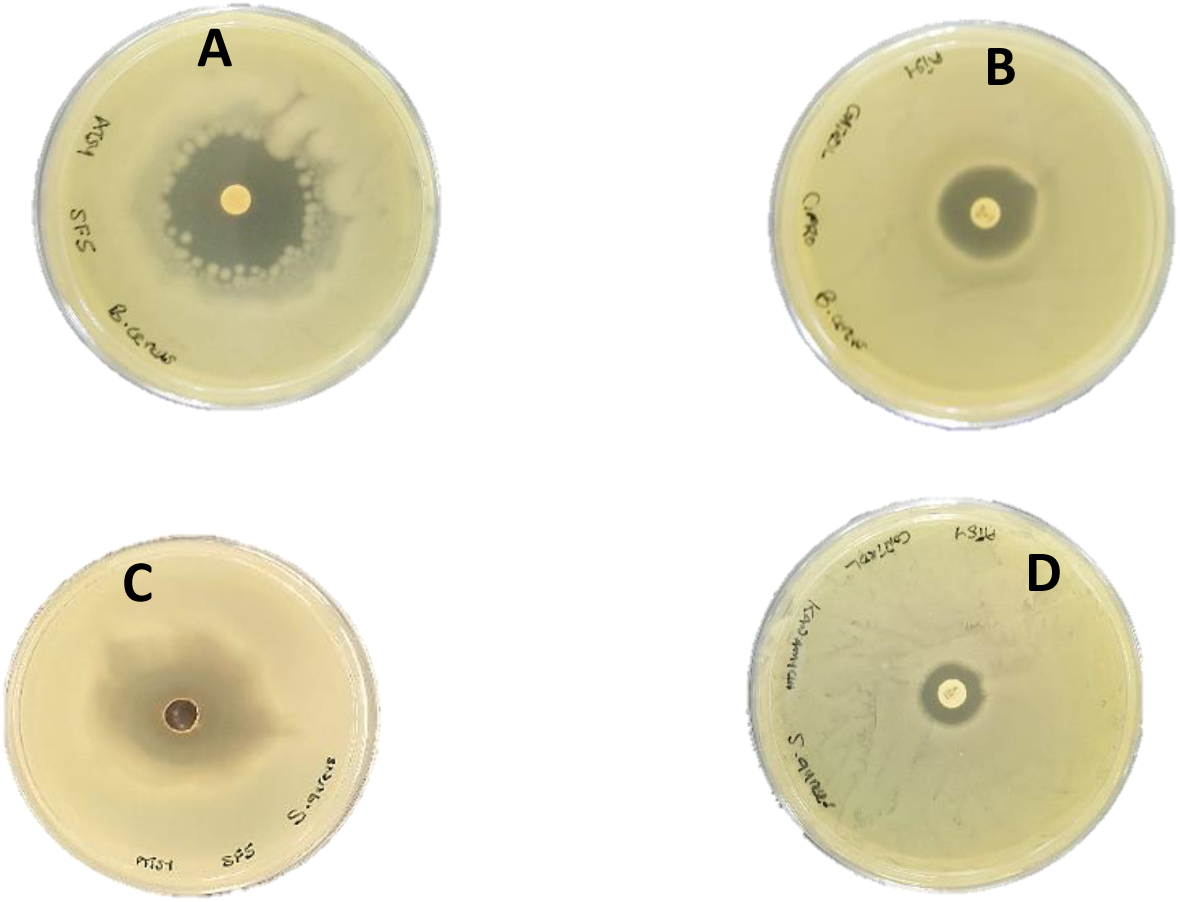
Zone of inhibitions shown by SF5 (10 mg mL^-1^), Ciprofloxacin (5 μg) and Kanamycin (30 μg) standard antibiotics against *B. cereus* ATCC 14579 and *S. aureus* ATCC 33591. (**A**) Zone of inhibition of **SF5** on *B. cereus*, (**B**) Zone of inhibition of Ciprofloxacin on *B. cereus*, (**C**) Zone of inhibition of **SF5** on *S. aureus* and (**D**) Zone of inhibition of Kanamycin on *S. aureus.*

Minimum inhibitory concentration (MIC) and minimum bactericidal concentration (MBC) was evaluated to ascertain the potency of the subfractions. The MIC and MBC values of the subfractions against the bacteria pathogens were 0.1 to 5 mg ml^-1^ (Table 3). The most susceptible pathogen to subfraction SF5 is *S. aureus.* The results indicated that SF5, with concentrations between 0.1 and 1 mg mL^-1^, could inhibit (bacteriostatic) and kill all the five pathogens (bactericidal) effectively. The three subfractions SF3, SF4 and SF6, with concentrations of 1 to 5 mg mL^-1^, were only able to inhibit (bacteriostatic) and kill (bactericidal) *B. cereus* and *K. pneumoniae*. In addition, 5 mg mL^-1^ of SF6 could only inhibit *K. pneumoniae* (bacteriostatic). Consequently, it is highly suggestive that SF5 comprise of the most active compounds against the five pathogens (Table 3).

**Table 3.** Minimum inhibitory concentration (MIC) and minimum bactericidal concentration (MBC) of Subfractions SF1-SF6 against five bacteria pathogens.

| Subfraction | SF3 (mg mL<sup>-1</sup>) |  | SF4 (mg mL<sup>-1</sup>) |  | SF5 (mg mL<sup>-1</sup>) |  | SF6 (mg mL<sup>-1</sup>) |  |
| --- | --- | --- | --- | --- | --- | --- | --- | --- |
| Organisms | MIC | MBC | MIC | MBC | MIC | MBC | MIC | MBC |
| *S. aureus* | 1 | 10 | 0.5 | 1 | 0.1 | 0.1 | 0.5 | 1 |
| *B. cereus* | 1 | 5 | 0.1 | 1 | 0.1 | 0.5 | 1 | 5 |
| *S. pyogenes* | 5 | - | 5 | - | 0.1 | 0.5 | 5 | - |
| *E. faecalis* | 5 | - | 5 | - | 0.1 | 0.5 | 5 | - |
| *K. pneumoniae* | 1 | 5 | 1 | 5 | 0.1 | 1 | 5 | - |

SF1-SF6 represent the sub-fractions from the second fractionation. Pathogens-*Pseudomonas aeruginosa* ATCC 10145, *Salmonella Typhimurium* ATCC 14028, *Proteus vulgaris* ATCC 8427, *Escherichia coli* ATCC 25922 and *Candida albicans* ATCC 90028 were resistance to all the fractions.

### Ultra-Performance Liquid Chromatography-Mass Spectrometry analysis (UPLC-MS)

The chromatogram of UPLC/MS analysis of the subfraction SF5 showed three series of ion peaks at [M+H]^+^ m/z = 197.1575 and 169.0823, at [M+H]^+^ m/z = 678.4408, at [M+H]^+^ m/z = 244.1696; 270.1950 and 270.2010 (Fig. 4A). In the first series, a set of mass peaks with an interval of 28 was observed and it has been reported that for the lipopeptides (depending on the type) a set of 14 or 28 molecular weight is often observed with different numbers of methylene (−CH_2_−) or ethylene (−CH_2_− CH_2_−) in fatty acyl chains [22 – 24]. The second series of peaks corresponds to a class of oligopeptide antibiotics such as monamycin (mol. wt. 677.84) and Alphostatin (mol. wt 668.637) produced by *B*. *megaterium* [25]. The third series of peaks with an interval of 26 molecular weight, cannot be immediately classified to compounds reported in the literature, though the mass data shows that it could be an ionic product of larger compounds. The first and second series of peaks in this study can be said to represent different fengycin and surfactin homologues by comparing their mass data with product ions obtained in previous studies [23, 26, 27].

**Figure 4.**
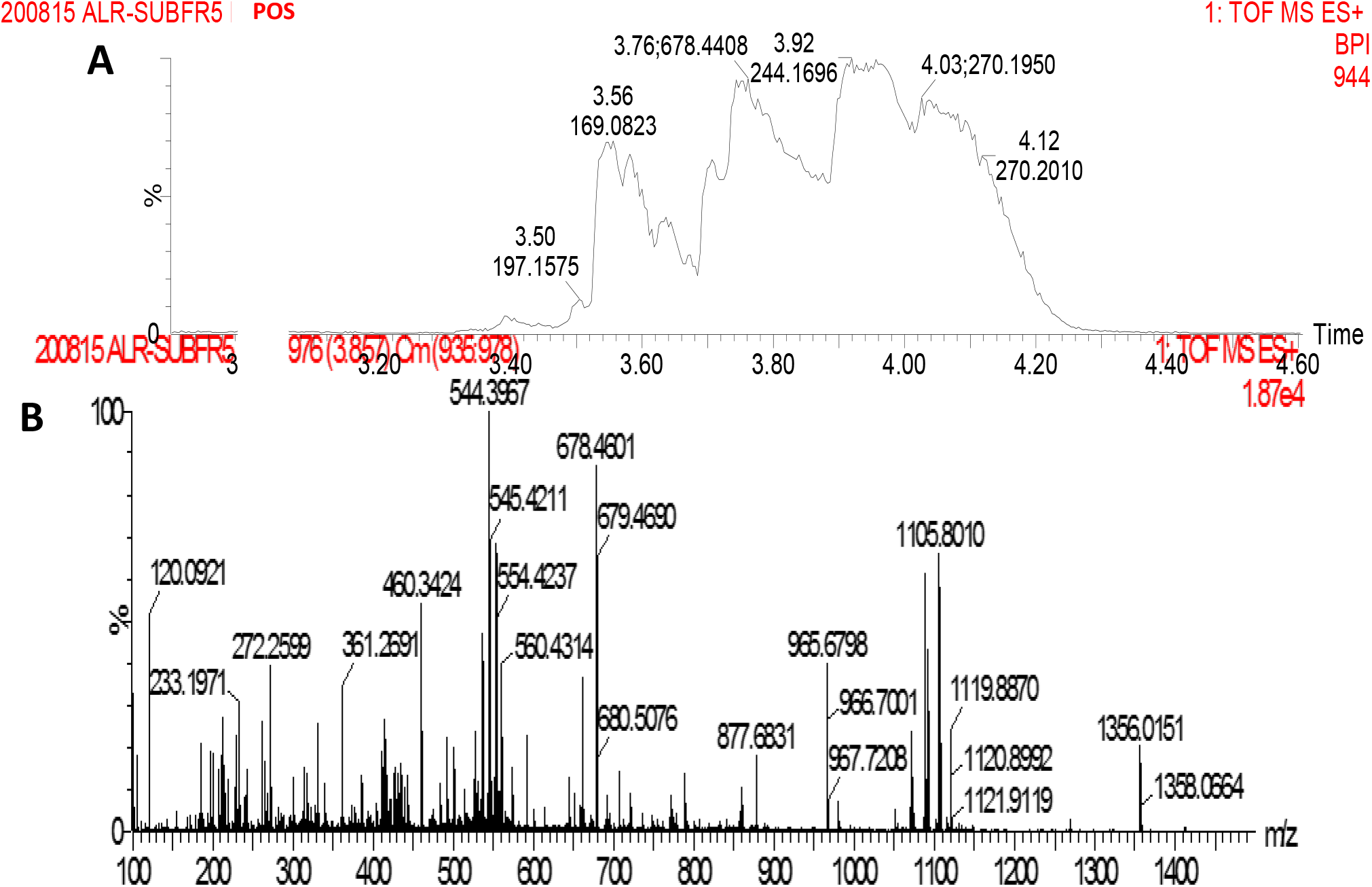
**(A)** Chromatogram of UPLC-MS of Subfraction SF5 for the time of flight (TOF) of 5 mins., showing the three series of ion peaks at [M+H]^+^ m/z = 197.1575; 169.0823, at [M+H]^+^ m/z = 678.4408, at [M+H]^+^ m/z = 244.1696; 270.1950; 270.2010 and **(B)** MS/MS spectra of precursor ions of m/z = 1356.0151; 1358.0664 and 1105.8010; 1119.8870; 1120.8992; 1121.9119 (Lipopeptides homologues).

The ion peak results showed product ions of [M+H]^+^m/z = 965.6798; 966.7001 and 967.7208, which corresponds to the collision ionization dissociation (CID) spectra of precursor ions of [M+H]^+^ m/z = 1356.0151 and 1358.0664 (Fig. 4B). The product ion fragments of [M+H]^+^ m/z = 965.6798; 966.7001 and 967.7208 can be said to be losses of fatty acid-Glu (amino acid residue) from the N-terminus segment of a lipopeptide antibiotic (fengycin B) [23, 27]. Similarly, the product ions of [M+H]^+^ m/z = 678.4601; 679.4690 and 680.5076 can be explained as ionized product of oligopeptide antibiotics, i.e. Monamycin (mol. wt. 677.84), as its accurate mass correspond to the [M+H]^+^ 678.4601, which has also been reported as one of the products of *Bacillus* sp. [17, 28]. The third series of peaks [M+H]^+^ m/z = 120.0921; 233.1971; 277.2599; 361.2691; 460.3424; 544.3967; 545.4211; 554.4237 and 560.4314 see Fig. 4B, can also be explained to be product ions from the precursor ions of [M+H]^+^ m/z =1105.8010; 1119.8870; 1120.8992 and 1121.9119 that belonged to fengycin isoforms [23].

The precursor ion [M+H]^+^ m/z = 1119.8870 observed, corresponded to Antibiotic complex 61-26 (mol. wt. 1120.348) produced by *Bacillus* sp. isolated by Wakisaka and Koizumi [29] and Shoji, Sakazaki [17]. *Bacillus* strains have been reported to produce several lipopeptide antibiotics and lipopeptide biosurfactants that have the capacity to inhibit Gram-positive bacteria and phytopathogenic fungi. They are so diverse that different strains of *Bacillus* produce different types of these substances [27, 30]. It is evident that endophytic *Bacillus tequilensis* ALR-2, has equal potential to produce series of lipopeptides antibiotics and fengycin isoforms which are responsible for its antimicrobial activities.

This study is also the first to report on the production of Antibiotic complex 61-26 (mol. wt. 1120.348) and Monamycin (mol. wt. 677.84) by the endophytic *Bacillus tequilensis* ALR-2. Conclusively, the UPLC-MS analysis has suggested that the precursor ions are lipopeptides (fengycins) and surfactins. Fengycins and surfactins are natural lipopeptides that show antimicrobial, antiviral and antitumor activities [23] and they are said to be safer in the environment when compared with other chemical agents. Fengycin are known to inhibit filamentous fungi but it does not inhibit yeast. In this study *Bacillus tequilensis* ALR-2 was found to be a co-producer of fengycins and surfactin as its antimicrobial activities and UPLC-MS have suggested. These natural lipopeptide compounds can be of use in the food preservation and control of plant diseases. Moreover, we suggest the determination of its amino acids and fatty acids residues to be able to elucidate its structure which may be useful in predicting it mechanism of action.

## Experimental procedures

### Culture establishment

The *Bacillus tequilensis* ALR-2 strain (KJ689792) was isolated from *A. vera* and identified as described in our previous study [9]. For this study, cultures were prepared on nutrient agar (Sigma-Aldrich) and incubated at incubation at 30 ± 2 °C for 18-24 h.

### Extraction of bioactive compounds

*Bacillus tequilensis* ALR-2 colony was inoculated into sterile Erlenmeyer flask (1L) containing 500 mL nutrient broth (Sigma-Aldrich) and incubated at 35 ± 2°C (150 rpm, 36 - 40 h) in Rotary Flask Shaker. The culture was centrifuged at 8,000 × *g* for 5 min to obtain the cell free supernatant. The extracellular metabolites in the cell free supernatant were exhaustively extracted with ethylacetate, diethylether and n-hexane separately in the ratio of 1:1 (supernatant:solvent). The organic phase was concentrated in rotatory evaporator (Rotavap) at 37°C, freeze-dried (LABCONCO freeZone 4.5^-105^°^C^) and weighed. The percentage yield of each extracts (% w/v) was calculated from the following equation: % extract yield = weight of dry extract (mg)/volume taken for extraction (1L) x 100

### Antimicrobial activity screening of the extracts via TLC-bioautography

The ethylacetate extract was fractionated on aluminium-backed thin layer chromatography (TLC) plates (Merck, silica gel 60 F254). The plates were developed under saturated conditions with the eluent systems, developed in our laboratory. The mobile phase used for separation of the bioactive metabolites was Chloroform:Methanol (4:1 v/v). The separations were detected under ultraviolet light (254 nm) and the Rf values were calculated. TLC-bioautography assay was carried out as described in our earlier publication [12] using immersion method. To develop the chromatograms, 1μL (20 mg mL^−1^) of the extract was loaded onto TLC plates in a narrow band and eluted using the mobile solvent system in a covered chromatography jar.

The developed plates were dried under the laminar flow for 6 h to remove traces of solvent on the TLC plates. One mL of 0.5 McFarland standard bacteria pathogens were mixed with molten Mueller Hinton agar in sterile petri dish. The prepared chromatograms were immersed in the molten Mueller Hinton agar and removed to solidify. This process was carried out in a laminar flow cabinet (Labotec, SA). The plates were then incubated in petri dish overnight at 37 ± 2°C and 100% relative humidity in the dark. The chromatograms were then sprayed with a 5 mg mL^-1^ solution of tetrazolium compound (TTC) and incubated overnight. Formation of white bands indicated bacterial growth suppression due to the presence of compounds that inhibited the growth of tested organisms that prevented the reduction of TTC to formazan (reddish coloration) [13].

### Extraction and purification of bioactive compounds

The crude ethylacetate extract of ALR-2 was purified via column chromatography using a LiChroprep RP-18 (15-25 μm) silica gel column (Merck, EMD Millipore Corporation, USA) and chloroform:methanol (ratio 4:1) as eluent. A total of 10 (5 mL each) fractions were collected and were lyophilized to obtain powder forms for the antimicrobial assay. Three fractions with good antimicrobial activities were subsequently combined and purified with acetonitrile:methanol as eluent to obtain a further six subfractions. The purity of each subfraction was analysed by reversed phase HPLC (COSMOSIL Packed C_18_ column, 4.6 ID × 150 mm, Agilent Technologies 1260 infinity). The following mobile phase was used; methanol:water with gradient elution method of 95% deionized water and 5% methanol for (0 – 20) min for solvent-A, and 5% deionized water and 95% methanol for (20 – 30) min for solvent-B.

### Antimicrobial assay

The antimicrobial activities of the six subfractions were assayed using the Kirby-Bauer disc diffusion method [14, 15], against bacteria and yeast pathogens, which *include-Pseudomonas aeruginosa* ATCC 10145*, Enterococcus faecalis* ATCC 29212*, Staphylococcus aureus* ATCC 33591*, Bacillus cereus* ATCC 14579*, Salmonella* Typhimurium ATCC 14028*, Proteus vulgaris*-ATCC 8427*, Klebsiella pneumoniae* ATCC 10031, *Escherichia coli* ATCC 25922*, Streptococcus pyogenes* ATCC and *Candida albicans* ATCC 90028. All pathogens were obtained from the Microbiology Laboratory of Monash University Malaysia. The bacteria pathogens were first pre-cultured overnight in Mueller Hinton broth (Difco, USA.) at 35 ± 2°C. Then, 5 mL of the culture were pipetted and centrifuged at 6,000 × *g* for 5 min. The pellets were re-suspended in sterile distilled water and the cell density was subsequently adjusted to 0.5 McFarland standard. The bacterial suspension was then seeded onto Mueller Hinton agar (Difco, USA.) plates for antimicrobial testing. The subfractions (10 μL of 10 mg mL^-1^) were impregnated onto sterile discs and placed on seeded agar plates. The plates were incubated at 35 ± 2°C for 48 h and the zone of inhibition was determined by measuring the diameter of annular clear zone. The experiment was performed in triplicates. Ciprofloxacin (5 μg) and Kanamycin (30 μg) standard antimicrobial discs were used as positive control.

### Minimum inhibitory concentration (MIC) and Minimum bactericidal concentration (MBC)

Microdilution method was used as described by Andrews [14]. The dilution range of the antimicrobial compound tested was modified to 0.1 mg mL^-1^ to 10 mg mL^-1^ concentrations. The bacterial inoculum was prepared and adjusted to 10^8^ cfu mL^-1^ (0.5 McFarland standards). Each antimicrobial extract (75 μL) was added into two rows of sterile 96-wells. This was followed with 75 μL of tested bacterial inoculum which was added into one row and 75 μL of control sterile broth was added into the second row of wells. In addition, inoculated and uninoculated wells of antimicrobial free broth were included to determine adequacy of the broth to support growth and to check for sterility of the broth. The microplate was covered with lid and incubated at 37 ± 2°C for 18 h. The absorbance of the culture turbidity was then read at 625 nm using the microplate reader (Infinite 200 TECAN). To assess MBC, wells with absence of growth (non-turbid wells) detected at the lowest concentration was sub-cultured onto Mueller Hinton agar (Difco, USA.) plates to determine the MBC. The concentration with no growth of tested organism is the MBC for the antimicrobial agent against the tested organism.

### Ultra-Performance Liquid Chromatography-Mass Spectrometry (UPLC-MS) analysis of antimicrobial compound

Ultra-Performance Liquid Chromatography-Mass Spectrometry (UPLC-MS) analysis was performed on the partially-purified antimicrobial substance using AcquityTM Waters Ultra Performance Liquid Chromatography (UPLC) (Waters Corporation, USA). The electrospray source was operated at a capillary voltage of 2.7kV. Column specification used was ACQUITY UPLC BEH C18 1.7 μm, 2.1 x 50 mm. Mass spectrometer used was Synapt high definition mass spectrometer quadrupole-orthogonal acceleration with time of flight detector (Waters Corporation, USA). Mobile phase used was: milliQ water + 0.1% formic acid (solvent A) and methanol + 0.1% formic acid (Solvent B). Flow rate was 0.5 mL/min. Gradient program was pre-determined as follows: 95-95% (solvent A) 0-2.8 min, 5-5% (solvent B) 0-2.8 min, 95-5% (solvent A) 2.8-3.5 min, 5-95% (solvent B), 5-95% (solvent A) 3.5-4.0 min 95-5% 3.5-4.0 min, 95-95% (solvent A) 4.0-5.0 min 5-5% (solvent B) 4.0-5.0 min.

## Statistical analysis

Results were expressed as mean ± Standard deviation (SD). One-way ANOVA was used to analyse all data obtained. The analysis was carried out using the Statistical Package for Social Science (SPSS) version 20.0 and means were compared using Tukey’s Studentized Range Test (HSD (0.05)), with differences considered statistically significant when p<0.05.

## Data availability

All data are contained in the manuscript. Accession number of organisms used are also indicated in the manuscript

## Acknowledgements

This work was supported by scholarship of Higher Degree for Research (HDR) School of Science, Monash University Malaysia.

## Conflict of interest

The authors report no conflict of interest and are responsible for the content and writing of the manuscript.

## References

[1]. Kobayashi D, Palumbo J. (2000). Bacterial endophytes and their effects on plants and uses in agriculture. Microbial endophytes. 2000:199–233.

[2]. Strobel GA. (2003). Endophytes as sources of bioactive products. Microbes and infection. 2003;5(6):535–44.

[3]. Sun L, Lu Z. (2005). Advance on antibiotics produced by endophytes. Food Ferment Ind (China). 2005;31:39–41.

[4]. Yu H, Zhang L, Li L, Zheng C, Guo L, Li W, et al. (2010). Recent developments and future prospects of antimicrobial metabolites produced by endophytes. Microbiological research. 2010;165(6):437–49.

[5]. Cho SJ, Park SR, Kim MK, Lim WJ, Ryu SK, An CL, et al. (2002). Endophytic Bacillus sp. isolated from the interior of balloon flower root. Bioscience, biotechnology, and biochemistry. 2002;66(6):1270–5.

[6]. Yang X, Strobel G, Stierle A, Hess W, Lee J, Clardy J. A (1994). fungal endophyte-tree relationship: Phoma sp. in Taxus wallachiana. Plant Science. 1994;102(1):1–9.

[7]. Huang J-S. (1986). Ultrastructure of bacterial penetration in plants. Annual review of phytopathology. 1986;24(1):141–57.

[8]. Guo B, Dai J-R, Ng S, Huang Y, Leong C, Ong W, et al. (2000). Cytonic acids A and B: novel tridepside inhibitors of hCMV protease from the endophytic fungus Cytonaema species. Journal of Natural Products. 2000;63(5):602–4.

[9]. Akinsanya MA, Goh JK, Lim SP, Ting ASY. (2015). Diversity, antimicrobial and antioxidant activities of culturable bacterial endophyte communities in Aloe vera. FEMS microbiology letters. 2015;362(23).

[10]. Gatson JW, Benz BF, (2006). Chandrasekaran C, Satomi M, Venkateswaran K, Hart ME. Bacillus tequilensis sp. nov., isolated from a 2000-year-old Mexican shaft-tomb, is closely related to Bacillus subtilis. International journal of systematic and evolutionary microbiology. 2006;56(7):1475–84.

[11]. Dawson J, Guilhaus M. (1989). Orthogonal-acceleration time-of-flight mass spectrometer. Rapid Communications in Mass Spectrometry. 1989;3(5):155–9.

[12]. Akinsanya, M. A, Ting, ASY, Sanusi, MO. (2017). Extraction methods and TLC-bioautography for evaluation of antimicrobial activities of endophytic bacteria from medicinal plants. LASU Journal of Medical Sciences, Volume 2(1), January-June 2017.

[13]. Begue, W. J., & Kline, R. M. (1972). The use of tetrazolium salts in bioauthographic procedures. Journal of Chromatography A, 64(1), 182–184.

[14]. Bauer A, Kirby W, Sherris JC, turck, Turck M. (1966). Antibiotic susceptibility testing by a standardized single disk method. American journal of clinical pathology. 1966;45(4):493.

[15]. Andrews JM. (2001). Determination of minimum inhibitory concentrations. Journal of antimicrobial Chemotherapy. 2001;48(suppl 1):5–16.

[16]. Suleiman MM, McGaw L, Naidoo V, Eloff J. (2010). Detection of antimicrobial compounds by bioautography of different extracts of leaves of selected South African tree species. African Journal of Traditional, Complementary and Alternative Medicines. 2010;7(1).

[17]. Shoji JI, Sakazaki R, Wakisaka Y, Koizumi K, Mayama M. (1975). Studies on antibiotics from the geneus *Bacillus.* V. Isolation of a new peptide antibiotic complex 61-26. The Journal of antibiotics. 1975;28(2):129–31.

[18]. Ruma K, Kumar S, Prakash H. (2013). Antioxidant, Anti-Inflammatory, Antimicrobial and Cytotoxic Properties of Fungal Endophytes from Garcinia Species. International Journal of Pharmacy and Pharmaceutical Sciences. 2013;5(3):889–97.

[19]. Sun L, Lu Z, Bie X, Lu F, Yang S. (2006). Isolation and characterization of a co-producer of fengycins and surfactins, endophytic Bacillus amyloliquefaciens ES-2, from Scutellaria baicalensis Georgi. World Journal of Microbiology and Biotechnology. 2006;22(12):1259–66.

[20]. Joseph B, Priya RM. (2011). Bioactive compounds from endophytes and their potential in pharmaceutical effect: a review. Am J Biochem Mol Biol. 2011;1(3):291–309.

[21]. Roongsawang N, Thaniyavarn J, Thaniyavarn S, Kameyama T, Haruki M, Imanaka T, et al. (2002). Isolation and characterization of a halotolerant Bacillus subtilis BBK-1 which produces three kinds of lipopeptides: bacillomycin L, plipastatin, and surfactin. Extremophiles. 2002;6(6):499–506.

[22]. Thaniyavarn J, Roongsawang N, Kameyama T, Haruki M, Imanaka T, Morikawa M, et al. (2003). Production and characterization of biosurfactants from Bacillus licheniformis F2. 2. Bioscience, biotechnology, and biochemistry. 2003;67(6):1239–44.

[23]. Kowall M, Vater J, Kluge B, Stein T, Franke P, Ziessow D. (1998). Separation and Characterization of Surfactin Isoforms Produced byBacillus subtilisOKB 105. Journal of colloid and interface science. 1998;204(1):1–8.

[24]. Morikawa M, Hirata Y, Imanaka T. (2000). A study on the structure–function relationship of lipopeptide biosurfactants. Biochimica et Biophysica Acta (BBA)-Molecular and Cell Biology of Lipids. 2000;1488(3):211–8.

[25]. Aoyagi T, Morishima H, Kojiri K, Yamamoto T, Kojima F, Nagaoka K, et al. (1989). Alphostatin, an inhibitor of alkaline phosphatase of bovine liver produced by Bacillus megaterium. J Antibiot (Tokyo). 1989;42(3):486–8. Epub 1989/03/01. PubMed PMID: 2496067.

[26]. Kim P, Bai H, Bai D, Chae H, Chung S, Kim Y, et al. (2004). Purification and characterization of a lipopeptide produced by Bacillus thuringiensis CMB26. Journal of applied microbiology. 2004;97(5):942–9.

[27]. Wang J, Liu J, Wang X, Yao J, Yu Z. (2004). Application of electrospray ionization mass spectrometry in rapid typing of fengycin homologues produced by Bacillus subtilis. Letters in applied microbiology. 2004;39(1):98–102.

[28]. Bevan K, Davies J, Hassall C, Morton R, Phillips D. (1971). Amino-acids and peptides. Part X. Characterisation of the monamycins, members of a new family of cyclodepsipeptide antibiotics. Journal of the Chemical Society C: Organic. 1971:514–22.

[29]. Wakisaka Y, Koizumi K. (1982). An enrichment isolation procedure for minor Bacillus populations. The Journal of antibiotics. 1982;35(4):450–7.

[30]. Pueyo MT, Bloch Jr C, Carmona-Ribeiro AM, Di Mascio P. (2009). Lipopeptides produced by a soil Bacillus megaterium strain. Microb Ecol. 2009;57(2):367–78.

